# S-Equol: A Novel Therapeutic For HIV-1-Associated Gastrointestinal Dysbiosis

**DOI:** 10.1101/2025.11.05.686823

**Authors:** Mason T. Rodriguez, Sarah J. Olmstead, Kristen A. McLaurin, Charles F. Mactutus, Rosemarie M. Booze

## Abstract

HIV-1 infection affects approximately 38.4 million people around the world. The advent of combination anti-retroviral therapy (cART) has greatly improved the quality of life of infected individuals; however, roughly 50% of these individuals will still experience HIV-1-associated neurocognitive disorders (HAND). Additionally, the gastrointestinal microbiome has been reported to be dysbiotic in HIV-1 infected individuals, regardless of adherence to cART. Current research has pointed to the gut-brain-microbiota axis as a potential target to treat both cognitive deficits and microbial changes. The present study investigated S-Equol (SE) as a potential therapeutic for HAND by modulating the gastrointestinal microbiome. The study included 21 HIV-1 Tg rats and 21 F344 control animals to test the effect 0.2mg SE has on cocaine-maintained responding on a PR schedule of reinforcement. Gastrointestinal microbiome alterations between genotypes were found at the phylum and genus level, regardless of treatment group, and SE treatment had both main effects and interactions with genotype. *Prevotella_UCG_001* was significantly associated with lever presses for drug, suggesting an effect on motivation for cocaine. *Alloprevotella* was found to significantly differentiate between genotype by treatment effects, indicating that SE differently affects genotypes.

## 1. Introduction

Human immunodeficiency virus 1 (HIV-1) has infected an estimated 38.4 million people around the world (1). Combination antiretroviral therapy (cART) is the main treatment for HIV and works to control the replication of HIV, resulting in improved CD4 T cell counts and preventing transmission by those compliant with the cART regimen to others (WHO, 2022). The use of cART has also greatly decreased the prevalence of HIV-1-associated dementia (HAD), but HIV-1-associated neurocognitive disorders (HAND) continue to persist, affecting approximately 50% of HIV-1 seropositive individuals regardless of cART treatment (2). Treatment is usually started as soon as a diagnosis is made but by then the infection has already established latent, viral reservoirs, that prevent full eradication of the virus by antiretrovirals, due to an inability of cART to reach these latent reservoirs (3). The infection enters the body through the exchange of bodily fluids (blood, semen, and vaginal secretions) with an infected individual. Once inside the body, HIV-1 initially binds to CD4+ T cells via the C-C chemokine receptor type 5 (CCR5) co-receptor. The gastrointestinal (GI) tract contains the largest mucosal immune system, making it one of the initial sites of HIV-1 infection. The GI tract is also one of the most damaged by the initial infection, indicated by a greater reduction in CD4+ T cells than in other tissues (4). After prolonged infection, the GI tract maintains the lowest level of CD4+ T cells, which is accompanied by alterations to the composition of the gastrointestinal microbiome (5,6,7,8).

It is well understood that dysbiosis, or negative alterations, of the gastrointestinal microbiome, can cause cognitive deficits through interactions with the gut-brain-microbiota axis (9). In HIV-1 seropositive individuals, dysbiosis of the gastrointestinal microbiome is seen throughout infection (6), with alterations in the microbiota composition found less than 6 months post-infection (8). HIV-1 infection results in an overall reduction in microbiota diversity, with a significant decrease in *Akkermansia Muciniphila* and an increase in *Prevotella* (6). Reduced *Akkermansia Muciniphila* and increased *Prevotella* is an important observation for two reasons: First *Akkermansia Muciniphila* is a vital bacterium that is responsible for maintaining the integrity of the gastrointestinal microbiome, and more specifically the epithelial barrier (10). Second, *Prevotella* is a gram-negative bacterium, meaning it possesses lipopolysaccharide (LPS) on the outer membrane, an endotoxin that increases immune activation and is commonly used to assess microbial translocation due to its ability to compromise the blood-brain barrier (BBB) (5,6,11). The combination of these two alterations allows HIV-1 to break down the beneficial bacteria in the gastrointestinal microbiome, compromising the integrity of the epithelial barrier in the process and opening the pathway for harmful microbes from the gastrointestinal tract to leak out. One of these harmful microbes is *Prevotella*, carrying LPS through the gut-brain-microbiota axis to the BBB where it simultaneously weakens the BBB as it passes through and binds to the surface of microglia (12,13). Changes in biomarkers of gastrointestinal epithelial barrier function or microbial translocation, such as circulating LPS, are correlated with immune dysfunction and were found to strongly predict mortality in HIV-1 seropositive individuals (10,14). Once LPS binds to the microglia, it puts them into an overactive state, damaging the cells over time and inadvertently causing the microglia to shed HIV-1 proteins from their latent reservoirs. Interestingly, elite controllers (people whose replication of HIV-1 is controlled without treatment) have microbiomes resembling uninfected individuals (3). Elite controllers are a unique example that illustrates how dysbiosis of the gastrointestinal tract could be influencing the progression of HAND. Many different infections and diseases have been associated with dysbiosis, indicating either the disease has altered the microbiome directly or it is the body’s response to compensate for the ongoing infection. There is a need to find a treatment that can simultaneously restore the gastrointestinal microbiota and prevent the over-activation of microglia, preventing the shedding of HIV-1 proteins into the brain.

The development of a treatment to prevent dysregulation of gastrointestinal microbiota has led to an increased interest in phytoestrogen compounds as a potential therapeutic for HAND. Phytoestrogens refer to plant-derived compounds that mimic mammalian estrogen and act on estrogen receptors (ERs). The main groupings of phytoestrogens are polyphenols, flavonoids, and isoflavonoids. These can be further broken down, with the most studied being lignans, flavonols, and isoflavones (15). Phytoestrogen compounds are found in most fruits and vegetables with the most abundant being in soybeans and other legumes (16). Phytoestrogen compounds were originally investigated as a possible supplement for women who were postmenopausal and had low estrogen levels (17,18). Phytoestrogen treatment was found to improve self-reported quality of life and cognitive symptoms among postmenopausal women. These initial successful findings on phytoestrogen treatments have led to an increased interest in phytoestrogens, particularly if these improvements would translate to men as well. Improvements in both psychomotor speed and spatial memory were found in two separate studies looking at adult men (19) and older, overweight adults (20) receiving isoflavone supplementation. Follow-up studies have found evidence for microglia inhibition as a mechanism by which phytoestrogens improve cognition and modulate brain activity (21, 22). To address the efficacy of phytoestrogens for treating symptoms of HAND, we investigated SE’s therapeutic effects in HIV-1 Transgenic (Tg) rats.

HIV-1 Tg rats express 7 of the 9 genes associated with HIV-1, providing a model for long-term HIV-1 viral protein exposure without active infection, resembling a state similar to HIV-1 positive individuals undergoing long-term cART therapy. Most, if not all, phytoestrogens require some metabolization by the gastrointestinal tract for there to be any health benefit, and a metabolite that is of particular interest is SE. It is generally ingested as Daidzein or Genistein and then broken down by gastrointestinal bacteria where it eventually ends up as SE, which is one of the metabolites important for the health benefits associated with phytoestrogen consumption (23). Individuals who metabolize daidzein into SE are equol producers and encompass ∼40% of individuals who eat a primarily Western diet with low soy (24).

Due to the possibility of the HIV-1 Tg rats not having the bacteria required to metabolize specific phytoestrogens, SE was selected as the treatment to bypass any confound that soy metabolization could cause. Additionally, the animals were ovariectomized to eliminate the potential confounding effects of endogenous estradiol. Estradiol is the natural estrogenic compound produced that can potentiate the reinforcing efficacy of cocaine (25), prevent HIV-1-induced neuronal damage (26,27,28), suppress HIV-1 transcription (29), and possibly affect the efficacy of SE treatment. The main goal was to investigate if SE protects or restores the microbiome from alterations associated with HIV-1. A secondary goal was to assess if these microbial changes in the gastrointestinal microbiome were associated with changes in motivation. Additionally, intestinal tissue samples were taken at the end of the experiment, these were used to assess differences in microbial composition between genotypes. The current hypothesis was that SE would alter responding to cocaine on a PR schedule of reinforcement by protecting against HIV-1-associated alterations in microbial composition. Specifically, HIV-1 will be differentiated by an increase in the bacteria *Prevotella* and a decrease in the bacteria *Akkermansia Muciniphila*. SE treatment will restore these bacterial levels in HIV-1 Tg rats to control levels, and these alterations will be associated with behavioral changes in the rats.

## 2. Materials and Methods

### 2.1 Animals

A total of 42 adult female ovariectomized (OVX) (21 HIV-1 Tg and 21 F344/N control) rats were purchased from Harlan Laboratories, Inc. (Indianapolis, IN, USA). A statistical power analysis and estimate of variance were performed to determine the number of animals needed for the proposed study. The power analysis suggests a sample size of n=20 per group to detect genotype (F344 vs. HIV-1 Tg) differences in specific bacteria (i.e., Prevotella). The sample size estimates should be adequate for determining significant differences with a power of 80% at an alpha level of 0.05. Another method of power analysis was performed that focused on gene sequencing reads and subject count; this power analysis estimate also suggested a sample size of n=20 per group would be adequate to determine significant differences with a power of 87% at an alpha level of 0.05.

All animals were ovariectomized at Harlan Laboratories; animals were ovariectomized due to the potential for estradiol effects to confound SE treatment effects. Animals were fed a low phytoestrogen diet (≤20 ppm of phytoestrogen; Teklad 2020X Global Rodent Diet; Harlan Laboratories, Inc., USA); standard rodent chow contains ∼350ppm of soy and alfalfa (Harlan Laboratories, Inc., IN). Both soy and alfalfa contain phytoestrogens, soy specifically contains daidzein which can be converted by gut bacteria to SE, which is the drug intervention used in the present study. All animals were maintained in an AAALAC-accredited facility using the guidelines established in the Guide for the Care and Use of Laboratory Animals of the National Institutes of Health. Animals had *ad libitum* access to food and water unless otherwise specified. The animal colony was maintained at 21±2°C, 50±10% relative humidity, and a 12L:12D cycle with lights on at 0700h. The Institutional Animal Care and Use Committee (IACUC) of the University of South Carolina approved the project protocol under federal assurance (#D16-00028).

### 2.2 Data Collection

HIV-1 Tg and control F344/N animals were trained to lever press for sucrose and then cocaine while being treated with either SE or sucrose pellets. A mixed-design ANOVA was used to analyze the impact SE and genotype had on responding behaviors using both a fixed and progressive ratio schedule of reinforcement. At the end, a choice behavior task was given to determine which reinforcer was preferred. Microbiome samples were collected at baseline and at the end of the study.

### 2.3 Drugs

Cocaine hydrochloride (Sigma-Aldrich Pharmaceuticals, St. Louis, MO) was weighed and dissolved in saline (0.9%). The solutions were made before the animals entered the operant chambers each day. Sucrose solutions were made fresh each testing day as well. SE (0.05mg) was purchased from Cayman Chemical (Ann Arbor, MI) and sucrose pellets (100mg) were purchased from Bio-Serv (Flemington, NJ). The SE pellets were added into 100mg sucrose pellets by Bio-Serv prior to sending them to the University of South Carolina, the combination of SE and the sucrose pellets was done to make sure both treatments were similar when administering SE or sucrose.

Additional drugs required for the use and maintenance of IV catheters included Heparin, purchased from APP Pharmaceuticals (Schaumburg, IL), Gentamicin sulfate from VEDCO (Saint Joseph, MO), butorphanol (Dolorex) from Merk Animal Health (Millsboro, DE), and Sevoflurane, USP from Baxter (Deerfield, IL).

### 2.4 Experimental Design

Animals were randomly assigned based on genotype to either SE (HIV-1-E = 11, F344/N-E=11) or sucrose (HIV-1-S=10, F344/N-S=10) treatment groups. Treatment groups received either 0.2mg SE (4 pellets) or 4 sucrose pellets per animal for 70 days. The dose of 0.2mg SE was selected for two reasons. First, it is consistent with what is used in human studies (∼20mg for ∼60kg human) and lower than the typical daily intake of phytoestrogens by elderly Japanese individuals (30-50 mg)(32). Second, using a dose-response experimental design, 0.2 mg SE was established as the most efficacious dose for the alleviation of sustained attention deficits in the HIV-1 Tg rat (33). Treatment started once daily for one week before the start of testing, and every day until catheterization. Following surgery, animals did not receive treatment for one week. Treatment resumed every other day until the end of the 14-day cocaine self-administration progressive ratio task. Treatment did not occur during the cocaine dose-response or choice behavior tasks. Animals were pair-housed for the duration of the sucrose tasks but were placed in separate cages to ensure the consumption of the pellets. After catheterization, the animals were single-housed and provided treatment in their home cage. Figure 1 provides an overview of the sequence followed.

**Figure 1.**
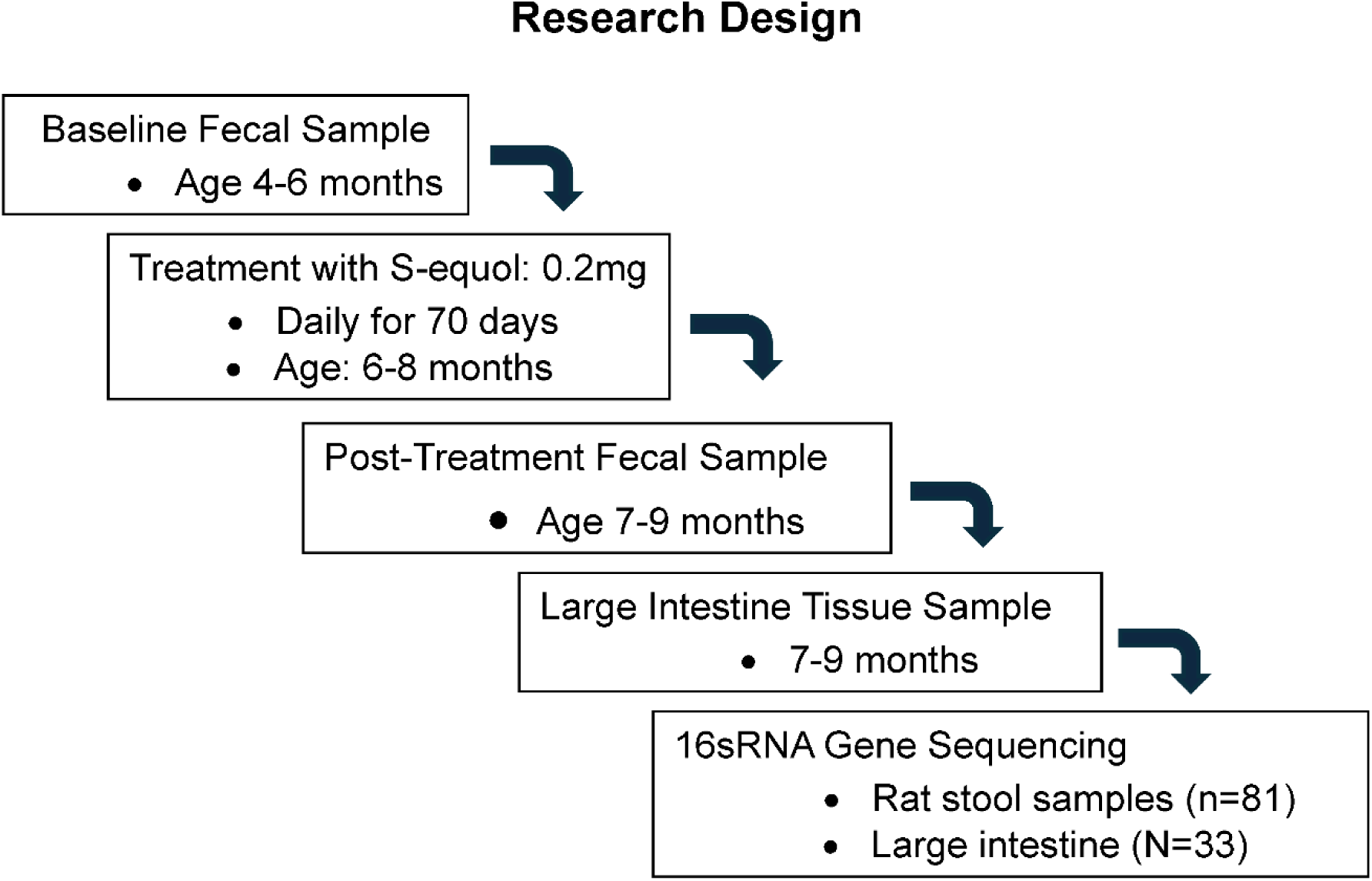
Research design illustrating the sequence employed to investigate if SE protects or restores the microbiome from alterations in the HIV-1 Tg rat.

### 2.5 Operant Chambers

Sound-attenuating enclosures housed operant chambers (ENV-008; Med Associated, St. Albans, VT) and were controlled by Med-PC computer software. Stainless steel was used for the front and back panels while the sides and top consisted of polycarbonate. The front stainless-steel panel contained a magazine that allowed a recessed 0.01cc dipper cup (ENV-202C) to deliver a solution through a 5cm x 5cm opening following the completion of a response requirement (ENV 202M-S). Two retractable active metal levers (ENV-112BM) on either side of the receptacle were located 7.3cm above the metal grid floor. The cue light was a 28-V white light, 3cm in diameter, and located above each active response lever but was never illuminated. Head entries into the magazine were detected using an infrared sensor (ENV 254-CB). There was another non-retractable lever located on the center of the back panel and a 28-V house light located above the lever. Responses on the center back lever were recorded but not reinforced. A syringe pump (PHM-100) was used to deliver intravenous cocaine infusions through a water-tight swivel (Instech 375/22ps 22GA; Instech Laboratories, Inc., Plymouth Meeting, PA), connected to the back mount of the animal using Tygon tubing (ID, 0.020 IN; OD, 0.060 IN) enclosed by a stainless-steel tether (Camcaths, Cambridgeshire, Great Britain). The infusion times of the pump were calculated by a Med-PC program according to the animal’s body weight (weighed daily).

### 2.6 16S rRNA Gene Sequencing

A total of 81 fecal samples and 33 intestinal tissue samples were collected and sent to the Alkek Center for Metagenomics and Microbiome Research (CMMR) at Baylor College of Medicine in Houston, Texas for the microbiome analysis. Samples were collected in sterile tubes and stored at - 80°C until being shipped. For shipping, samples were placed in order and shipped on dry ice overnight with a sample manifest that included de-identified sample IDs and the tube positions. The analysis pipeline for the gene sequencing uses custom packages created by the CMMR to provide summary statistics, quality control measurements, multi-run reports, and characterization of microbial communities across large numbers of samples. In brief, DNA was extracted from fecal samples using PowerMag Soil DNA Isolation Kit (MoBIO Laboratories, CA) and intestinal tissue samples using Power Lyser Ultra Clean Tissue and Cell RNA kit (MoBio Laboratories, CA) both according to the manufacturer’s protocol. 16S rRNA gene sequencing was done using the V4 primer region on the Illumina MiSeq program (Illumina, CA) to generate a baseline and the microbiome’s response to SE. Gene sequences were clustered into operational taxonomic units (OTUs) based on the 16Sv4 region, and phylogenetic, alpha, and beta-diversity changes were all reported.

### 2.7 Cocaine-Mediated Responding

Animals were first trained on an FR1 schedule of reinforcement (0.2mg/kg/inj) for 5 consecutive days, each session lasting 1 hour. Following completion of the response, the requirement resulted in a 20s time-out where the animals could not respond. In the next phase of the project, animals responded for IV cocaine on a PR schedule of reinforcement (0.75mg/kg/inj) for 14 consecutive days, each session lasting a maximum of 120 minutes. Completion of each ratio resulted in a 20s time-out.

### 2.8 Data Analysis

Data analysis was performed using SPSS (IBM Corporation, Armonk, NY) and Graphpad (Graphpad Software, Inc., La Jolla, CA). A 2x2 factorial design was used to analyze the bacterial changes of the microbiome with genotype (HIV-1 Tg vs. F344/N control) and treatment (SE vs. Sucrose) as between-subject factors. Diversity measures were calculated at baseline and after completion of treatment to determine pre- and post-changes from SE and if there was an interaction between genotype and treatment group. The alpha diversity analysis examined the bacterial richness and evenness within samples, which included OTUs (richness), Chao1 (estimator of diversity), and Shannon Diversity Index (richness and evenness). Beta diversity was assessed via unweighted (dissimilarity based on phylogenetic differences but not taxonomic abundance) and weighted UniFrac analyses (dissimilarity based on phylogenetic differences and taxonomic abundance). A principal coordinates analysis (PCoA) approach was used to summarize the compositional differences of each microbiome sample. A follow-up MANOVA was done with SE treatment (SE vs. Sucrose) and genotype (HIV-1 Tg vs. F344/N control) as independent variables and Cocaine slope (escalation rate) and *Prevotella_UCG_001* change over time (difference between baseline and follow-up measure) as the dependent variables. Lastly, a discriminate function analysis was done to determine if the bacterial differences in Prevotella_UCG_001, *Alloprevotella*, and *Akkermansia muciniphila* could be used to discriminate groups based on genotype and treatment received. Animals with potential patency issues (back mount leakage or inability to flush) were excluded from the analysis. Significant differences were set at p≤0.05.

## 3. Results

### 3.1 16S rRNA Gene Sequencing

The alpha diversity analysis of baseline (stool) samples suggested some evidence for differences between HIV-1 Tg and F344/N control rats based on observed operational taxonomic units (OTUs; richness)(p≤0.051) but neither the estimates of diversity (Chao1) or richness and evenness (Shannon diversity index) even approached conventional levels of statistical significance (**Fig. 2A**). PCoA was performed using the unweighted and weighted UniFrac analysis of the beta diversity of baseline (stool) sample differences between genotypes. The unweighted UniFrac approach (dissimilarity based on phylogenetic differences) accounted for approximately 20% of the variance (r^2^=0.199) and was statistically significant at baseline (p≤0.001)(**Fig. 2B**). In contrast, the weighted UniFrac approach (dissimilarity based on phylogenetic differences and taxonomic abundance) accounted for less than 4% of of the variance (r^2^=0.037) and was not statistically significant (p≤0.21).

**Figure 2.**
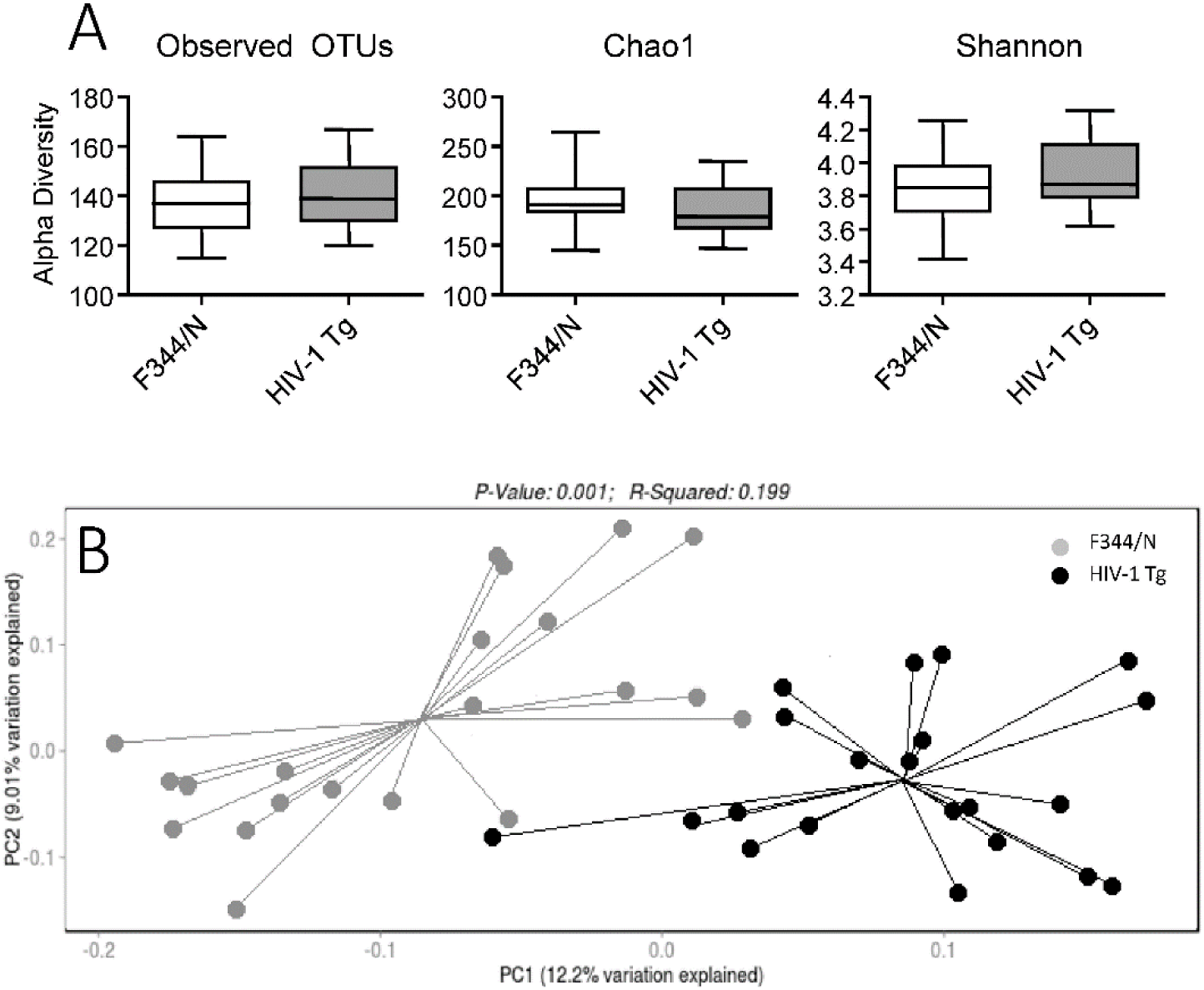
(A) Baseline differences in alpha diversity based on Observed operational taxonomic units (OTUs) neared significance (p≤0.051), still indicating no difference in taxonomy between genotypes. Chao1 (p≤0.2) and Shannon Index (p≤0.22) estimate the overall diversity and richness, respectively. Both were non-significant at baseline between genotypes. **(B)** Baseline beta diversity was presented via principle coordinates analysis using unweighted UniFrac measures of baseline bacterial composition, with significant differences being found at baseline measures between genotypes (p≤0.001).

With regard to tissue samples, the alpha diversity analysis of baseline (stool) samples suggested little evidence for differences between HIV-1 Tg and F344/N control rats based on observed operational taxonomic units (OTUs; richness)(p≤0.37), diversity (Chao1, p≤0.37), or richness and evenness (Shannon diversity index, p≤0.60). The assessment of Beta diversity using the unweighted UniFrac approach only accounted for 4% of the variance (r^2^=0.040) and was not statistically significant (p≤0.296). Similarly, the weighted UniFrac approach Beta diversity for tissue samples also only accounted for 4% of the variance (r^2^=0.040) and was not statistically significant (p≤0.28).

Alpha diversity measures of stool before and after treatment indicated no significant difference between HIV-1 Tg animals (p≤0.464) or F344/N controls (p≤0.235) based on observed OTUs, Chao1 (ps≤0.404) or of the Shannon diversity index (p≤0.481, p≤0.597, respectively)(**Fig. 3A**). In contrast, the results for the Beta diversity measures of stool indicated consistent positive effects of Se treatment. The unweighted UniFrac approach demonstrated a statistically significant effect of treatment for both the control F344/N (p<0.022; r^2^=0.057) and HIV-1 Tg animals (p≤0.035; r^2^=0.050), indicating a shift in phylogenetic makeup in both genotypes after SE treatment (Fig. 1D). Similarly, the results for the weighted UniFrac approach also displayed a statistically significantly effect of treatment for both the control F344/N (p<0.047; r^2^=0.063) and HIV-1 Tg animals (p<0.007; r^2^=0.101), indicating a shift in phylogenetic makeup and taxonomic abundance in both genotypes after SE treatment (**Fig. 3B**).

**Figure 3.**
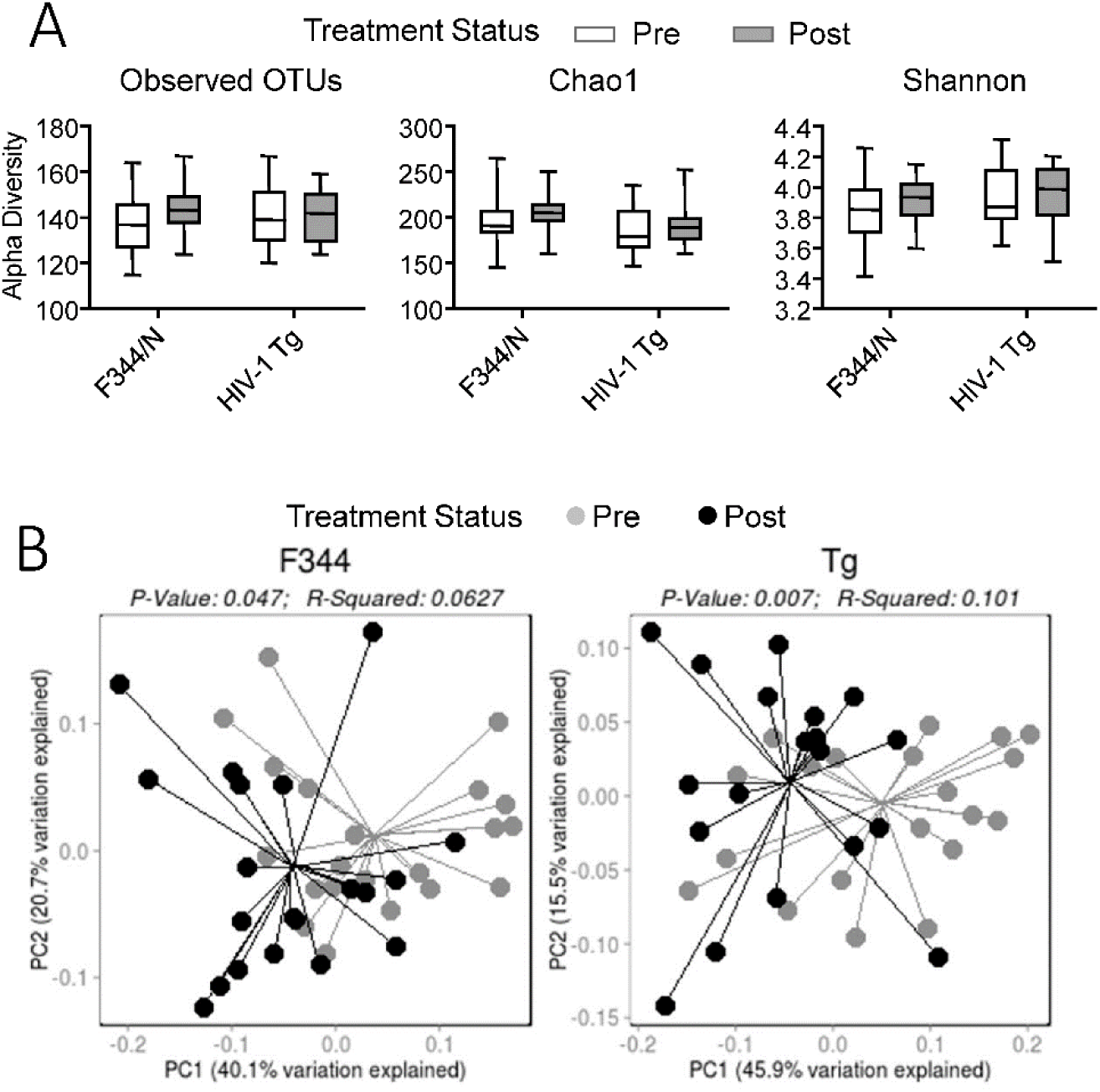
(A) Follow-up alpha diversity stool samples were analyzed to reveal treatment effects between genotypes. F344/N animals’ bacterial composition did not significantly differ after treatment, indicated by non-significant OTUs (p≤0.235), Chao1 (p≤0.404), and Shannon Index (p≤0.481). Similarly, HIV-1 Tg animals did not alter in alpha diversity after treatment, OTUs (p≤0.464), Chao1 (p≤0.404), and Shannon Index (p≤0.597). **(B)** Principal coordinates analysis (PCoA) of beta diversity change after treatment was done using weighted UniFrac measures of bacterial composition. Results indicate significant alterations in both F344 (p≤0.047) and HIV-1 Tg animals (p≤0.007) after treatment, suggesting SE affects both genotypes’ microbiome composition.

### 3.2 Microbiome Alterations

The bacterial makeup of the gastrointestinal microbiome was analyzed at the phylum and genus levels. Baseline samples were found to be non-significant between genotypes at the phylum level. There were, however, baseline differences at the genus level between genotypes when looking at the top 20 abundant bacteria. *Bacteroides*, *Alloprevotella*, *Streptococcus*, *Lachnoclostridium*, and *Tyzzerella* were all significantly elevated in HIV-1 Tg animals at baseline compared to F344/N control animals (**Fig. 4A**).

**Figure 4.**
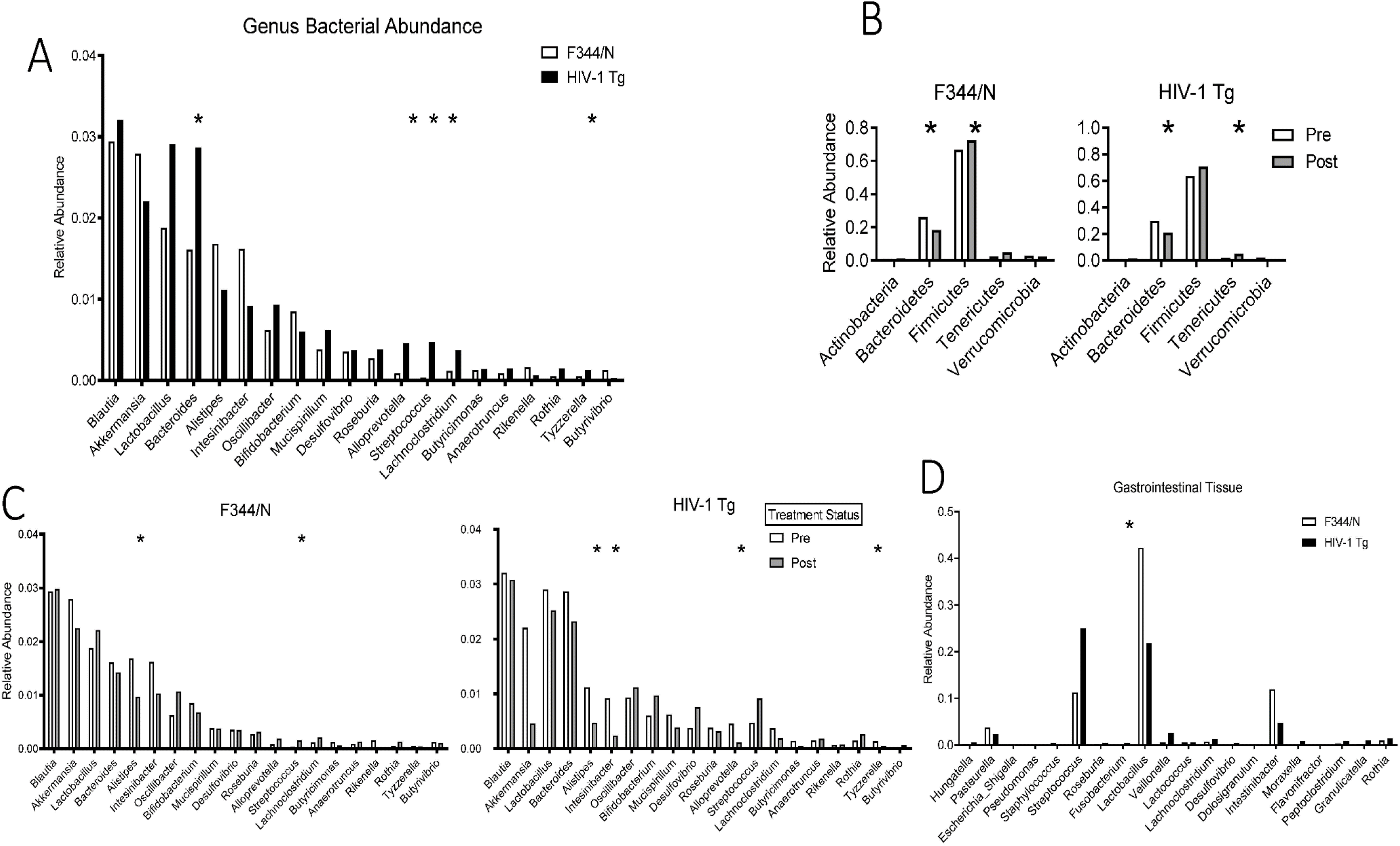
(A) Baseline genus-level taxonomic abundance was summarized according to the relative abundance of the top 20 genera from both genotypes. Significant baseline differences were found with HIV-1 Tg animals having increases in *Bacteroides*, *Alloprevotella*, *Streptococcus*, *Lachnoclostridium*, and *Tyzzerella* with alpha at 0.05. **(B)** Phylum-level significant alterations were present in *Bacteroidetes* and *Tenericutes*. The change in *Bacteroidetes* and *Tenericutes* was in both genotypes, with *Bacteroidetes* being lower and *Tenericutes* being higher after treatment. **(C)** Significant genus-level alterations in *Alistipes* were in both genotypes after treatment, both resulting in a reduction after treatment. Genotype x treatment differences were found with *Streptococcus* being increased in F344 animals while *Intestinbacter*, *Alloprevotella*, and *Tyzzerella* were reduced in the HIV-1 Tg animals, all at an alpha of 0.05. **(D)** Genus level differences in tissue samples were found with *Fusobacterium* increased in HIV-1 Tg animals compared to the F344 animals at an alpha level of 0.05.

SE treatment had a main effect at the phylum level by significantly decreasing *Bacteroidetes* and increasing *Tenericutes* in both genotypes (**Fig. 4B**). Additionally, at the genus level, SE treatment significantly lowered *Alistipes* in both genotypes. Within genotypes, F344/N control animals treated with SE experienced a significant increase in *Streptococcus* while HIV-1 Tg animals had a significant increase in *Intestinbacter*. HIV-1 Tg animals treated with SE also had significant decreases in *Alloprevotella* and *Tyzzerella* (**Fig. 4C**). Tissue samples were non-significant at the phylum level but at the genus level, *Fusobacterium* was significantly higher in HIV-1 Tg animals compared to F344/N control animals (**Fig. 4D**).

### 3.3 Prevotella Change with Cocaine Use in S-Equol-Treated Animals

Specific bacteria were chosen based on the relevance they have to HIV-1 status. The MANOVA analysis indicated an overall effect of genotype, F(3, 28)=3.358, p≤0.050 on the cocaine use slope and *Prevotella_UCG_001* change over time. When looking at specific between-subjects effects there was a main effect of genotype on *Prevotella_UCG_001* change over time, F(3, 28)=6.822, p≤0.014 but not Cocaine use slope, F(3, 28)=0.651, p≤0.426. Follow-up pairwise comparisons with Bonferroni corrections found that *Prevotella_UCG_001* change over time was significantly different between SE-treated F344/N and HIV-1 Tg animals, p≤0.027, 95% C.I.=[.001,.018] (**Fig. 5**) but no difference between Sucrose treated F344/N and HIV-1 Tg animals (p≤0.198).

**Figure 5.**
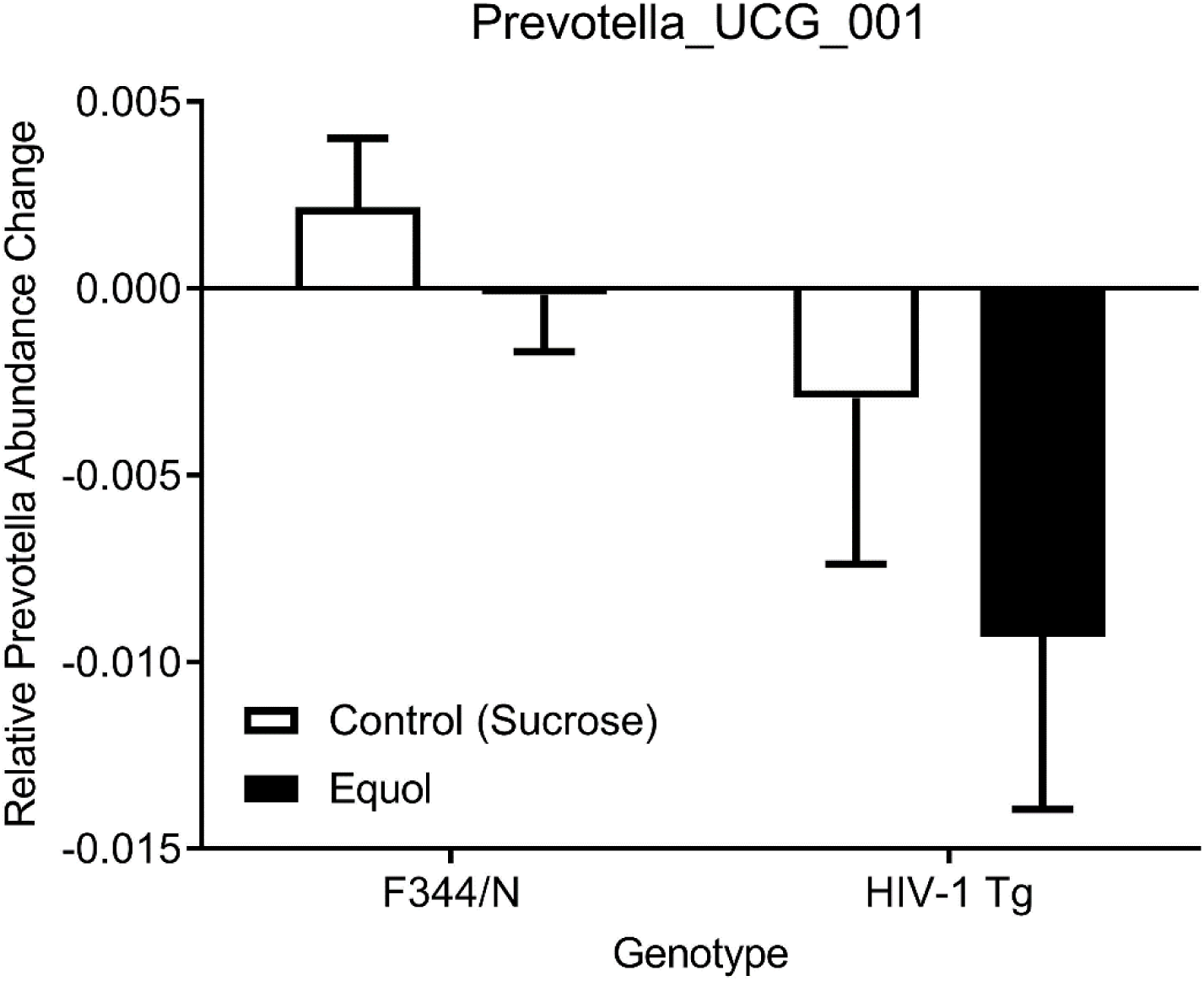
MANOVA analysis was done to determine if there was a difference in both *Prevotella_UCG_001* change over time and lever presses for cocaine based on genotype and treatment. An overall significant effect of genotype was found (p≤0.50) leading to a follow-up pairwise comparison with Bonferroni corrections that suggested the significant *Prevotella_UCG_001* change over time was due to the SE-treated F344/N and HIV-1 Tg animals (p≤0.027) as there was no difference in Sucrose treated animals (p≤0.198).

### 3.4 Bacterial Discriminants of Group Membership

A discriminate function analysis was performed to determine the best bacteria to differentiate between genotype x groups. A discriminant function analysis of the baseline bacteria revealed *Prevotella_UCG_001* and *Alloprevotella* to be significant, (λ=0.453, X^2^ = 21.782, R^2^ = 0.504, p≤0.010), with 56.3% of original grouped cases classified correctly. The significant discriminant function analysis indicates a difference in their abundance of *Prevotella_UCG_001* and *Alloprevotella* between genotype x treatment groups at baseline.

When investigating the post-treatment samples, a discriminant analysis successfully demonstrated the separation of genotype x treatment groups. Alloprevotella was found to significantly discriminate genotype x treatment effects, at an alpha of 0.05. The prominent bacterial change observed in HIV-1 Tg animals was attributed to Akkermansia displaying a large decrease in the equol treated HIV-1 Tg group, an effect not observed in the F344 controls administered equol (**Fig. 6**).

**Figure 6.**
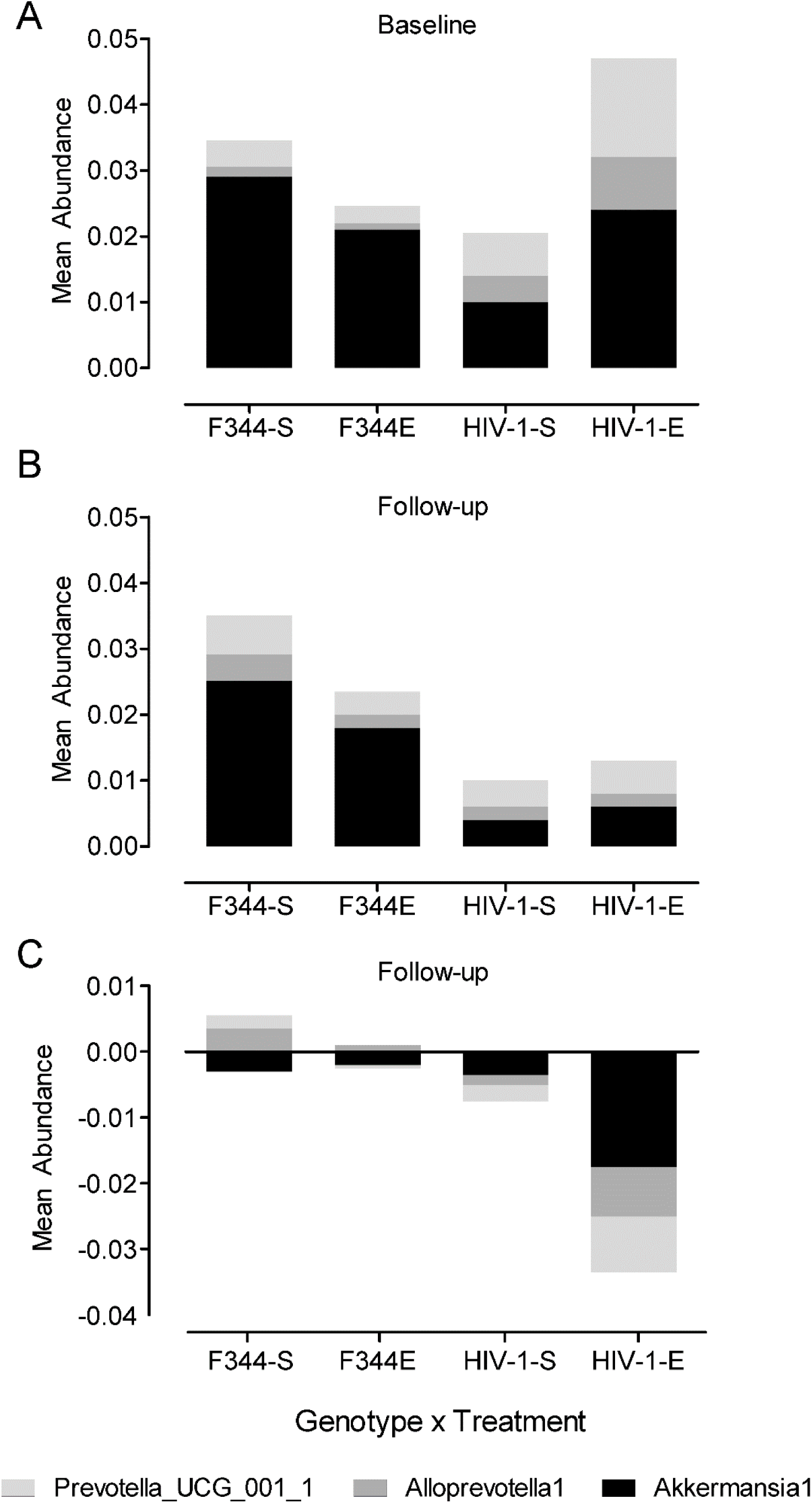
Discriminant function analysis was performed to determine if HIV-1-relevant bacteria could be used to separate genotype x treatment groups. At baseline (A), Prevotella_UCG_001 and Alloprevotella were significantly different between genotypes x treatment groups (p≤0.010). At follow-up (B), discriminant analysis of abundance differences in *Prevotella_UCG_001*, *Alloprevotella*, or *Akkermansia Muciniphila* confirmed no genotype x treatment effect. A prominent genotype effect was suggested with *Akkermansia Muciniphila* decreased due to HIV-1 Tg status, but SE treatment did not alter this change. In the bottom panel (C), discriminant analysis successfully demonstrated the separation of genotype x treatment groups at the end of the study. Alloprevotella was found to significantly discriminate genotype x treatment effects, at an alpha of 0.05. The prominent bacterial change observed in HIV-1 Tg animals was attributed to Akkermansia displaying a large decrease in the equol-treated HIV-1 Tg group, an effect not observed in the F344 controls administered equol.

## 4. Discussion

The gastrointestinal microbiome of HIV-1 seropositive individuals is characterized by an overall reduction in microbiota diversity, and at the genus level, a significant reduction in *Akkermansia Muciniphila* and an increase in *Prevotella* when compared to healthy individuals (5,6). In the present study, HIV-1 Tg rats were found to have a similar increase with a higher abundance of *Prevotella_UCG_001* in the gastrointestinal microbiome, but *Akkermansia Muciniphila* was non-significantly different even though there appears to be a reduction occurring based on Figure 4C. Previous studies have reported that even with cART, long-term HIV-1 infection leads to a significant loss of *Akkermansia Muciniphila* compared to healthy individuals (4,5,34). At the end of the study, a discriminant analysis successfully demonstrated the separation of genotype x treatment groups with the prominent bacterial change observed in HIV-1 Tg animals attributed to *Akkermansia Muciniphila* displaying a large decrease in the SE-treated HIV-1 Tg group, an effect not observed in the F344 controls administered SE. The reduction in *Akkermansia Muciniphila* ultimately is an important finding as it suggests that the HIV-1 Tg rat could be a useful model for studying a state of gastrointestinal dysbiosis similar to that found in HIV-1 seropositive humans.

HIV-1 Tg rats exhibit neurocognitive deficits in prepulse inhibition, learning, and sustained and selective attention and are associated with synaptodendritic alterations of medium spiny neurons (35–39). The current study highlights that in addition to these deficits, there are changes in the gastrointestinal microbiome composition between HIV-1 Tg rats and F344 control animals that could be influencing these deficits through interactions in the gut-brain-microbiota axis. HIV-1 Tg rats were found to have elevated *Prevotella_UCG_001* levels, a bacteria that possesses LPS on its outer membrane, an endotoxin that can damage the epithelial barrier and weaken the BBB (5,11). Increased LPS could be one of the main mechanisms that HIV-1 Tg-associated gastrointestinal dysbiosis is worsening symptoms of HAND. Once LPS passes the BBB, it can bind to the surface of microglia, increasing immune activation and putting the cell in an overactive state (12,13).

The current study investigated whether the HIV-1 transgene and SE treatment altered specific bacteria in combination with lever pressing for cocaine on a PR schedule of reinforcement that lasted 14 days. *Prevotella_UCG_001*, a subset of *Prevotella*, and lever pressing for cocaine were significantly different with genotype and treatment as independent variables. The change in *Prevotella_UCG_001* was investigated further and found to be dependent on genotype with HIV-1 Tg animals experiencing a larger change in *Prevotella_UCG_001*. Additionally, there was a difference between genotypes treated with SE, suggesting that the difference in *Prevotella_UCG_001* between genotypes could be attributed to SE treatment. The study also sought to determine if any of the specific bacteria could be used to differentiate the effects of genotype and treatment on lever pressing. The discriminant function analysis suggested that *Prevotella_UCG_001* and *Alloprevotella* could be used to differentiate genotype x treatment groups at baseline, with HIV-1 Tg rats possessing higher abundances of both *Prevotella_UCG_001* and *Alloprevotella*. Following treatment with SE, the discriminant function analysis revealed that *Prevotella_UCG_001* and *Alloprevotella* were indistinguishable between the genotype x treatment groups, indicating that both bacterial abundances of HIV-1 Tg rats had returned to similar levels as F344/N controls.

Together these findings indicate that SE can modulate the gastrointestinal microbiome composition, and these alterations have a direct link to the motivational changes being observed in the animals. It also reiterates that phytoestrogens appear to interact differently with ill vs. healthy animals, meaning that the use of phytoestrogens for a cognitive or behavioral treatment may not benefit everyone the same (40,41). For individuals who are already experiencing a deficit in cognition, phytoestrogens may be restorative, but if it is a healthy individual then phytoestrogen treatment may not have any effect.

As mentioned previously, SE is another compound that has also modulated motivational behaviors, specifically regarding cocaine intake. HIV-1 Tg rats who experience HIV induced deficits in motivational behaviors, indicated by a reduction in the reinforcing efficacy of sucrose compared to controls who escalated intake of sucrose during a sucrose maintained responding task, utilizing a PR schedule of reinforcement (42). The dysfunction was largely indicated by the findings associated with cocaine intake, which found that HIV-1 Tg rats escalated the intake of cocaine on a PR schedule, more so than the control animals, indicating a dysfunction of motivational behaviors as a natural reinforcer such as sucrose does not maintain an escalation pattern of responding but a drug reinforcer such as cocaine does increase in this manner. SE was found to restore the dysfunctional motivational responses by the HIV-1 Tg rats to cocaine (42).

The improvement in motivational dysfunction observed in HIV-1 Tg rats may be due to modulation of DAT. Evidence for DAT modulation has been reported (43) with Genistein administered to aging rats to investigate the impact it had on executive function and dopaminergic activity. Genistein is a phytoestrogen compound that preludes SE; when Genistein is ingested, depending on if the individual has the necessary bacteria, will metabolize Genistein into what eventually becomes SE (24). Chronic treatment with genistein increases the expression of DAT in old rats, resembling similar levels of those found in similar aged control animals (43). The present finding in combination with the work of others (42) indicate that prolonged treatment with phytoestrogen compounds could improve overall DAT function in HIV infected individuals (43).

Phytoestrogen treatment appears to improve motivational deficits, particularly in regard in substance use disorder and in the presence of HIV, in a multitude of ways. Supplementation with Phytoestrogens would improve the overall bacterial composition of the gut microbiome, reducing levels of systemic inflammation (44) and thereby reducing the circulation of LPS and the microbial translocation of infected monocytes (13). Treatment with phytoestrogens would also improve DAT functioning, which is already readily targeted by the HIV-1 tat protein and has been found to inhibit DAT (45,46). Another method that phytoestrogen supplementation would improve DAT functioning is through binding to microglia cells, reducing the number of overactivated microglia by binding the cells and preventing subsequent activation by LPS that binds to the microglial cells (47,48,49). The combination of these events suggests that there are many potential points to target the motivational dysfunction found in HIV infected individuals. The specific targets of phytoestrogens also seem to prevent or reduce HIV specific damage done to the CNS in addition to attenuating dysfunctions in reward-seeking behaviors (50).

Limitations to the present study are that sample collections were done only as pre and post-samples, therefore we cannot dissociate the effects of SE from cocaine on the microbiome. Ideally, a sample collection between the onset of SE and cocaine would allow an understanding of how SE affected the microbiome composition before the start of cocaine. Additionally, the antibiotic Gentamicin was administered IV following the implantation of the indwelling catheter. Notably, this antibiotic treatment was administered intravenously to all rats as a standard practice to maintain catheter patency. Although oral antibiotic treatment alters microbiome composition, there were no instances of oral antibiotic treatment during the study.

Overall, SE possesses large therapeutic potential for HIV-1-associated gastrointestinal dysbiosis by modulating *Prevotella_UCG_001* and *Alloprevotella* towards an abundance similar to the control animals. Reducing *Prevotella* would lead to less surrounding LPS circulating in the gastrointestinal tract and therefore lower the potential escape of LPS to the blood and the BBB. The bacterial alterations indicated an interaction between SE and the genotype, supporting the need for follow-up analysis on the efficacy of phytoestrogen use, specifically in diseased states. Further investigations also need to be done to discover the specific mechanism of action that SE uses to (1), alter the microbiome composition, and, (2), modulate neurocognition and motivated behavior. It will also be important to address, (3), why an individual’s state of health seems to modulate SE’s efficacy in regard to microbiotic gastrointestinal health and neurocognition.

## 5. Conclusions and Future Directions

The current study investigated the effect SE has on both cocaine-maintained responding and gastrointestinal microbiome composition in HIV-1 Tg rats and F344 control animals. The study found specific bacteria to be associated with lever pressing for cocaine and alterations strong enough to allow for accurate classification of group membership. Taken together, there is an important interaction between the HIV-1 transgene and the gastrointestinal microbiome, with specific bacterial differences similar to human individuals living with HIV-1. The gut-brain-microbiota axis has been reported to influence many behavioral and cognitive functions, suggesting the importance of improving dysbiotic states that go beyond possible local damage to the gastrointestinal tract but to wide arching effects.

Phytoestrogen compounds such as SE have gained much interest in recent years for their efficacy in improving cognition and well-being in individuals. In regard to HIV-1, SE and other phytoestrogens have the potential to migrate from the gastrointestinal microbiome to the brain where they can bind to microglia via estrogen receptors and g-protein-coupled receptors, with the most common one being GPR30 (22,26). Suggestions have been made as to how phytoestrogens can improve cognitive function and most include inflammation and/or microglia activation. Phytoestrogens bind to microglia to reduce overactivation and inflammatory markers, but how the reduction occurs is still up for debate.

Many different phytoestrogens have been discussed as possible candidates for adjunctive therapy to cART for the prevention of HAND. Resveratrol and Quercetin are the most studied compounds but Genistein and Daidzein are arguably the most important due to one of their main metabolites being SE. Much of the studies have found similar results with all phytoestrogens, with some overlap but some distinct pathways being used between the compounds. An important finding among all the phytoestrogen compounds was that they are only beneficial to individuals with deficits(51). Additionally, most studies found greater benefits when treatment was started early on than later, this was especially true with regards to HIV-1; a similar outcome is noted with SE in the HIV-1 Tg rat (33,38,39). Furthermore, phytoestrogens are important in treating HIV-1 because they are capable of binding to microglia via ERs, and more specifically GPR30 (22). LPS binds to microglia via TLR4 to exert their over activation, yet phytoestrogens such as Genistein have been found to prevent this over activation (see Fig. 7) and prevent damage to the cells (52). Phytoestrogens have also been found to improve gut health by increasing microbiota diversity and more importantly, has been found to reduce the level of LPS from a high fat diet (53). This points to phytoestrogens as a possible treatment for both gut dysbiosis and neurocognitive deficits related to HIV-1.

**Figure 7.**
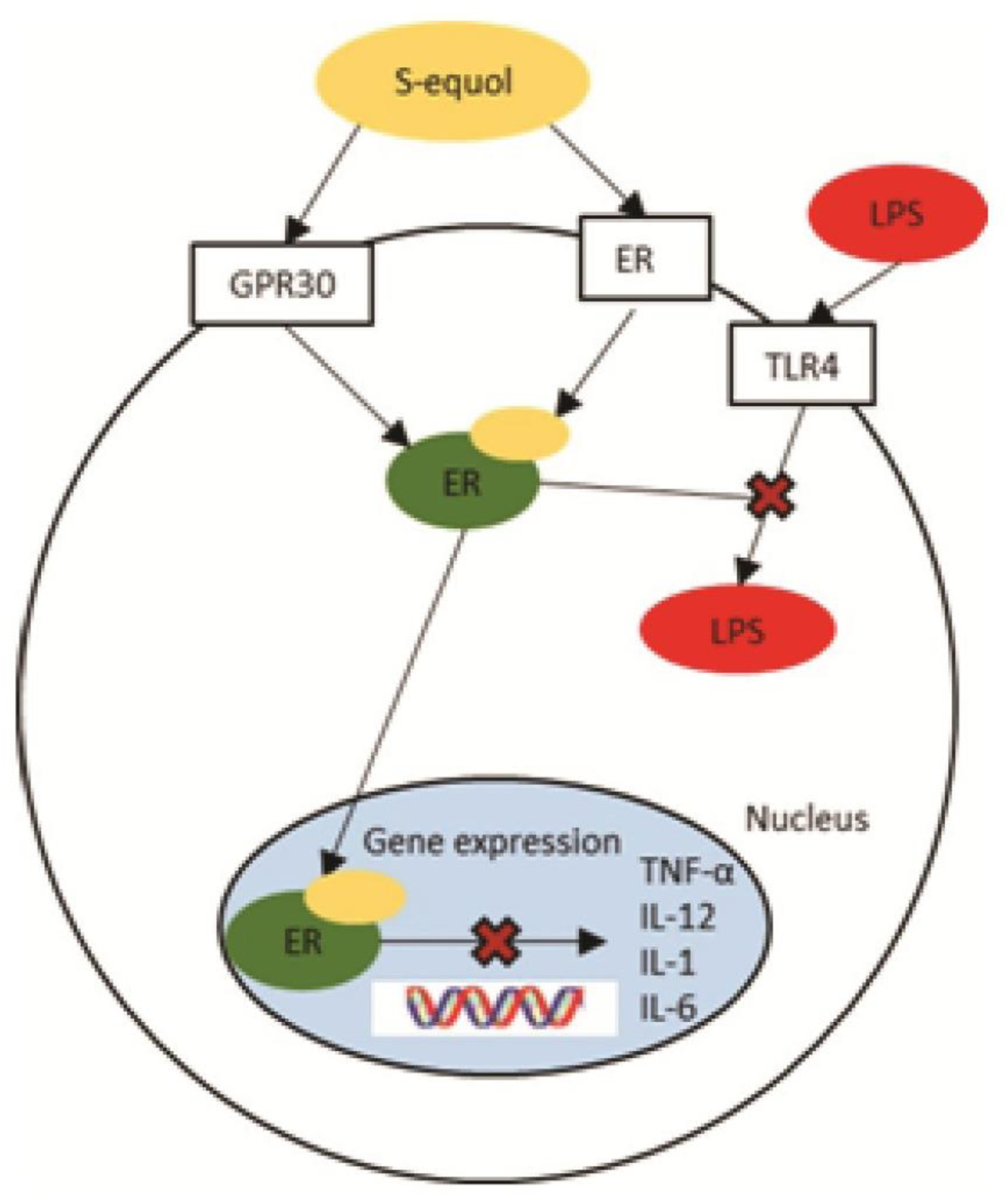
Schematic model of S-Equol and LPS Binding to Microglia. S-Equol travels via the gut-brain-microbiota axis into the CNS, where it binds to microglia via GPR30 and ERs. LPS also binds to microglia but via the TLR4 receptor. S-Equol prevents chronic activation via inhibition of LPS and prevents expression of inflammatory markers.

Limitations of the review are that many studies have not investigated the effects phytoestrogens have specifically on HAND and HIV-1 induced gut dysbiosis. Most studies have focused on either postmenopausal cognitive changes or age-related changes. More recently they have been investigated for their effects on diseases such as Alzheimer’s and dementia, but results must be interpreted and translated to the HIV-1 populations to understand if they have potential to treat HAND in conjunction with cART. Additionally, the mechanism by which phytoestrogens are able to prevent LPS induced over activation of microglia is still not fully understood. Previous studies have pointed to an interaction between ERs, G protein-coupled receptors, and toll-like receptors. Through binding to ERs or G protein-coupled receptors, phytoestrogens prevent over activation of microglia, either by lowering activation of the cell or somehow preventing the mechanism that LPS uses to over activate the cell. Further investigations need to be done to discover the mechanism of action which phytoestrogens prevent LPS-induced over activation of microglia.

Taken all together, phytoestrogens represent an innovative treatment addition to cART for the treatment of HAND and HIV-1-induced gut dysbiosis. HIV-1 has largely changed from a terminal illness to a chronic one that is characterized by neurocognitive deficits as age increases. Approximately 50% of seropositive individuals will experience HAND, illustrating the need for a treatment that can effectively and safely prevent these deficits from occurring. Alterations to the gut microbiome are well understood now to affect cognitive function and overall health, making the gut-brain-microbiota axis a potential treatment pathway that could alleviate not only the cognitive deficits related to HAND but also the dysbiotic state induced by HIV-1 infection.

Potential future directions include a follow-up cell culture study to investigate the effects of *Prevotella*-bound LPS on microglial function, then subject the cells to SE treatment, allowing for an investigation into how SE can directly modulate microglial function and by what mechanisms SE’s effect occurs. Having a greater understanding of SE mechanism may allow for more optimal HIV-1 treatment paradigms as HAND continues to affect approximately 50% of seropositive individuals, regardless of adherence to cART. Thus, SE may provide a novel adjuvant treatment in addition to cART for HIV-1-associated dysbiosis and resultant neurocognitive dysfunction.

## 6. Research Funding

This work was supported in part by grants from the NIH (National Institute on Drug Abuse, DA059310 to R.M.B. and C.F.M., and DA056288 to K.A.M.; National Institute of Neurological Disorders and Stroke, NS100624 to C.F.M. and R.M.B.; National Institute on Aging, AG082539 to C.F.M. and R.M.B.) and the interdisciplinary research training program supported by the University of South Carolina Behavioral-Biomedical Interface Program (GM081740). The sequencing was performed at the Alkek Center for Metagenomics and Microbiome Research at Baylor College of Medicine.

## 7. Ethical Approval

Animal care and use were performed in accordance with the Guide for the Care and Use of Laboratory Animals of the National Institutes of Health and were approved by the Institutional Animal Care and Use Committee of the University of South Carolina under federal assurance (#D16-00028).

## 8. Author Contributions

All authors have accepted responsibility for the entire content of this manuscript and approved its submission. Conceptualization, R.M.B., and C.F.M.; Data Collection, S.J.O.; Data Analysis, M.T.R., and C.F.M.; Writing—Original Draft Preparation, M.T.R.; Review and Editing, R.M.B., C.F.M., and K.A.M.; Funding Acquisition, R.M.B., C.F.M., and K.A.M. All authors have read and agreed to the published version of the manuscript.

## 9. Data Availability

The datasets generated and/or analyzed during the current study are available in the NCBI SRA repository, accession number is: PRJNA1088843

## 10. Competing Interests

The authors declare that they have no conflict of interest.

**Abbreviations**
HAND: HIV-1-associated neurocognitive disorders,
cART: combination anti-retroviral therapy
CCR5: C-C chemokine receptor type 5,
HIV-1 Tg rat: HIV-1 Transgenic rat,
SE: S-Equol,
BBB: blood-brain barrier,
PR: Progressive Ratio;
FR: Fixed Ratio

